# Classification and characterisation of brain network changes in chronic back pain: a multicenter study

**DOI:** 10.1101/223446

**Authors:** Hiroaki Mano, Gopal Kotecha, Kenji Leibnitz, Takashi Matsubara, Aya Nakae, Nicholas Shenker, Masahiko Shibata, Valerie Voon, Wako Yoshida, Michael Lee, Toshio Yanagida, Mitsuo Kawato, Maria Rosa, Ben Seymour

## Abstract

Chronic pain is a common and often disabling condition, and is thought to involve a combination of peripheral and central neurobiological factors. However, the extent and nature of changes in the brain is poorly understood. Here, we investigated brain network architecture using resting state fMRI data collected from chronic back pain patients in UK and Japan (41 patients, 56 controls). Using a machine learning approach (support vector machine), we found that brain network patterns reliably classified chronic pain patients in a third, independent open data set with an accuracy of 63%, whilst 68% was attained in cross validation of all data. We then developed a deep learning classifier using a conditional variational autoencoder, which also yield yielded 63% generalisation and 68% cross-validation accuracy. Given the existence of reliable network changes, we next studied the graph topology of the network, and found consistent evidence of hub disruption based on clustering and betweenness centrality of brain nodes in pain patients. To examine this in more detail, we developed a multislice modularity algorithm to identify a consensus pattern of modular reorganisation of brain nodes across the entire data set. This revealed evidence of significant changes in the modular identity of several brain regions, most notably including broad regions of bilateral sensorimotor cortex, subregions of which also contributed to classifier performance. These results provide evidence of consistent and characteristic brain network changes in chronic pain, and highlight extensive reorganisaton of the network architecture of sensorimotor cortex.

## Introduction

Maladaptive brain processing of pain is thought to have a primary or facilitative role in many types of chronic pain. In chronic back pain, for example, degenerative musculoskeletal change is considered unlikely in itself to fully explain persistent pain in most patients, and central processes are thought to be critical for the chronification and maintenance of pain. Existing data have identified a broad array of structural and functional brain differences in patients (Baliki, Geha, et al., 2008; Tagliazucchi et al., 2010; Baliki, Baria, and Apkarian, 2011; Kutch et al., 2017; Hemington et al., 2016; Napadow et al., 2010; Hashmi, Baliki, et al., 2013), and this has led to the concept of chronic pain as a brain network disorder (Apkarian, Baliki, and Geha, 2009; Mano and Seymour, 2015; Kuner and Flor, 2017). However, given the complexity of brain networks, we still do not have a reliable and consistent characterisation of these changes.

One of the difficulties in identifying robust changes in brain networks underlying chronic pain is that networks are inherently data-rich, and the patterns of disruption may be complex. One way to tackle this is to use machine learning and deep learning methods, and a number of studies have shown how this can be used to successfully build biomarkers (i.e. classifiers) in a range of psychiatric disease (Yahata et al., 2016; Takagi et al., 2017; Watanabe et al., 2017; Yamada et al., 2017). However, these methods need to be validated on genuinely independent data sets to be convincing, and current evidence of generalisable classifiers for chronic pain is lacking.

Even so, interpreting brain network changes based purely on classifiers alone can be difficult. This is because the classifier pattern itself is often comprised of a large matrix of individual functional connections, and strongly predictive (i.e. information-rich) functional connections do not necessarily imply an active role in a disease. A better way of describing and understanding networks is to instead evaluate the underlying topology (Bullmore and Sporns, 2009; Bressler and Menon, 2010). Since the brain is inherently modular, individual differences in function can be reflected by differences in a number of network characteristics (Meunier, Lambiotte, and Bullmore, 2010). This approach offers a way to define specific aspects of network architecture that change in a disease.

With these issues in mind the aim of the current study was to i) to classify, and ii) characterise brain networks in chronic back pain in a multi-site study using resting state fMRI. For classification, we applied machine learning and deep learning classifiers based on data from two sites (Cambridge, UK and Osaka, Japan) as a discovery cohort, and used an open data set (Chicago, USA) as a validation cohort. For characterisation, we investigated hub disruption across all datasets, and developed a method to identify brain regions that undergo modular reorganisation in the chronic pain state.

## Methods

### Participants

Across two sites (UK and Japan), a total of 41 adults with chronic musculoskeletal low back pain (CLBP) and 56 approximately age, sex, and IQ-matched adults without CLBP participated in the study. Patients were recruited under the following inclusion criteria: Chronic back pain for over 6 months, no other chronic pain condition, no other major neurological or psychiatric disease, and no contraindications to MRI scanning. The study was approved by the local research ethics committee at Addenbrookes Hospital in Cambridge, and Osaka University Medical hospital in Osaka. Prior to the participation, all participants gave written informed consent.

For all participants, the pain scores were taken in the form of visual analog scale (VAS) and Short-Form McGill Pain Questionnaire. Mood information was collected with Beck Depression Inventory (BDI) and Hamilton Depression Rating Scores. IQ information was collected using the National Adult Reading Test (NART) 1 for the participants in the U.K., and the Japanese Adult Reading Test (JART) 2 for the participants in Japan. Their demographic information is summarised in Table 1.

**Table 1:** Demographic details of participants.

|  |  | *N* | Age | BDI | Duration | VAS | JART/NART |
| --- | --- | --- | --- | --- | --- | --- | --- |
| CLBP | JP | 24 | 21-66 | 15.2 ± 10.5 | 11.6 ± 9.2 | 2.6 ± 2.4 | 31.23 ± 9.25 |
|  | UK | 17 | 20-61 | 15.9 ± 11.5 | 10.4 ± 7.5 | 4.8 ± 2.8 | 29.31 ± 6.76 |
|  | US | 34 | 21-62 | 6.3 ± 5.8 | 15.7 ± 11.3 | 6.7 ± 1.7 | — — |
| TD | JP | 39 | 21-68 | 4.7 ± 3.4 | 0 | 0.3 ± 1.1 | 34.66 ± 7.38 |
|  | UK | 17 | 20-62 | 3.7 ± 5.3 | 2.4 ± 7.5 | 0.3 ± 0.7 | 37.29 ± 6.84 |
|  | US | 34 | 21-64 | 1.5 ± 2.6 | 0 | 0 | — — |

### MRI data acquisition

All the scans were performed on a 3.0-T MRI Scanner (3T Magnetom Trio with TIM system; Siemens, Erlangen, Germany) equipped with echo planar imaging (EPI) capability and a standard 12-channel phased array head coil either at Addenbrooke’s hospital (Cambridge, UK) or CiNet (Osaka, Japan). Participants remained supine and wore MR-compatible headphones with their heads immobililised with cushioned supports during scanning. Resting-state functional MRI (rsfMRI) was acquired using a single-shot EPI gradient echo T2*-weighted pulse sequence with the following parameters: for the participants in the U.K.; TR 2000 ms, TE = 30 ms, FA = 78 degrees, BW = 2442 Hz, FOV = 192 × 192 mm (covering the whole brain), acquisition matrix = 64 × 64, 32 axial slices with a interleaved slice order of 3.0mm slice thickness with 0.75mm inter-slice gap, 300 volumes; for the participants in Japan; TR 2500 ms, TE = 30 ms, FA = 80 degrees, BW = 2367 Hz, FOV = 212 × 212 mm (covering the whole brain), acquisition matrix = 64 × 64, 41 axial slices with a ascending slice order of 3.2mm slice thickness with 0.8mm inter-slice gap, 234 volumes. A high-resolution three-dimensional volumetric acquisition of a T1-weighted structural MRI scan was collected using a MPRAGE pulse sequence: for the participants in the U.K.; TR = 2300 ms, TE = 2.98 ms, time of inversion = 900 ms, FA = 9 degrees, BW = 240 Hz, FOV = 256 × 256 mm, 176 sagittal slices of 1mm slice thickness with no inter-slice gap, acquisition matrix = 256 × 256. for the participants in Japan; TR = 2250 ms, TE = 3.06 ms, time of inversion = 900 ms, FA = 9 degrees, BW = 230 Hz, FOV = 256 × 256 mm, 208 sagittal slices of 1mm slice thickness with no inter-slice gap, acquisition matrix = 256 × 256.

### Resting-state fMRI data preprocessing

High-resolution T1-weighted anatomical imaging and a resting-state functional imaging were performed for each participant, and all those images were preprocessed with SPM8 (Wellcome Trust Centre for Neuroimaging, University College London, UK) on Matlab (R2014a, Mathworks, USA). The first five volumes were discarded to allow for T1 equilibration. Slice timing was adjusted to the intermediate slice and all images were realigned to the first volume of each scan with the estimated 6 rigid-body head motion parameters. After T1 weighted structural image was co-registered to the mean EPI volume, tissue segmentation of the structural image into three tissue classes; gray matter (GM), white matter (WM), and cerebrospinal fluid (CSF), based on the T1-weighted image contrast was performed in the common Montreal Neurological Institute (MNI) space. The relevant parameters estimated in the tissue segmentation were applied to warp functional images into MNI152 template space with a 2 × 2 × 2 mm spatial resolution. Subsequently, smoothing was applied with a 6 × 6 × 6 mm FWHM Gaussian kernel.

### Inter-regional correlation analysis

To investigate the inter-regional functional relationship among regions over the whole brain, we used the digital BSA-AAL composite atlas composed of 140 ROIs consisting of the BrainVISA Sulci Atlas (BSA) and the Anatomical Automatic Labeling (AAL) package (with a spatial resampling of 2 × 2× 2 mm^3^ grid in MNI space); a band-pass filter with a transmission range from 0.008 to 0.1 Hz; regression out of the nuisance regressors from a mask of white matter, cerebrospinal fluid, and the whole brain based on the segmentation of individual T1 weighted image, and three translational and three rotational head motion parameters; To protect against motion artifact, we performed scrubbing with a threshold of frame displacement (FD) of 0.5 mm. This preserved 91.7% and 85.0% of slices in control and CLBP patients in the Japan data respectively, 81.3% and 75.4% in the UK data, and 93.5% and 88.8% in the US data. Subsequently a 140 × 140 Pearson’s full correlation matrix was computed on all pairs of each of intra-regional average time-series of the ROIs.

### Classification

A classification model built from UK and Japan data sets was tested on an open data set available from the “Open Pain Project” (http://www.openpain.org/, Department of Physiology, Northwestern University). Anatomical MRI data in the test data set were provided with skull stripping during preprocessing. We chose to exclude six participants (three CLBP patients and three controls had lost a small part of brain coverage in their anatomical image during the skull stripping procedure. The US dataset differed in only 1 aspect of the inclusion criteria (choosing to exclude patients with a BDI score of over 19).

### Classification using Support Vector Machines

We used a support vector machine (SVM) classifier (Cortes and Vapnik, 1995) based on the connectivity (correlation) matrices to classify subjects as patient or control. SVMs learn a hyperplane, or decision boundary, that separates the two classes as well as possible (i.e. maximises the margin between the samples in the two classes). Once this boundary is learnt new samples are classified according to the side of the hyperplane they fall. The optimal margin is parameterised by a weight vector, W. Each entry of W corresponds to a particular feature, in this case a connectivity measure between two brain regions, and is interpreted as the contribution of the feature to separating the classes. However, it is important to note that the predictions are based on all features. Linear kernel SVMs have only one hyperparameter, *C*, controlling the trade-off between the width of the margin separating the two classes and the number of misclassified samples.

To assess the predictive performance of the SVM classifier we ran two validation models: i) pooling together the UK and Japan data as the training dataset, and using the US data as an independent validation dataset *(validation model* 1); ii) pooling the three datasets together (UK, Japan and US) to increase power and testing the performance of the classifier using a stratified Leave-Two-Subjects-Out (LTSO) cross-validation (CV) *(validation model* 2). LTSO CV allowed for one subject of each class to be left out for testing, and the remaining subjects from both classes to be used for training in each CV fold. To account for multi-site effects, the pairs that were left out were always from the same acquisition site (stratification).

Due to the slightly higher number of controls compared to patients, we also bootstrapped 100 models in both validation approaches. In other words, each time we ran the whole validation model we randomly selected a balanced sample (as large as possible) with an equal number of patients and controls. The predictive accuracies were averaged across bootstraps.

Feature selection was carried out on the training data using a univariate two-sample t-test (Guyon and Elisseeff, 2003): we Fisher-transformed the correlation data and kept only the features (connections) statistically significant between patients and controls (*p* < 0.05, uncorrected).

For both validation approaches the SVM *C* parameter was optimised using grid search (between 10^−3^ to 10^3^) and a Leave-3-Out CV on the training data. This CV was nested within the LTSO CV in the second validation model.

We used as performance measures the accuracy (percentage of correctly classified samples) and both sensitivity and specificity (percentage of correctly classified patients and controls, respectively). The obtained results were tested for statistically significance (i.e. how unlikely the results would be if the classifier was randomly attributing the class labels) using a permutation approach, where we repeat the entire classification procedure (including the two validation models, parameter optimisation, bootstrapping and feature selection) 1,000 times, each time permuting the labels (patient or control) (Nichols and Holmes, 2002).

We used *python* 2.7.12 and the *scikit-learn* 0.17.1 machine learning library (Pedregosa et al., 2011) for this analysis.

### Classification using Deep Learning

We used a conditional variational autoencoder (CVAE) based on the 140 ROIs to classify subjects (Kingma et al., 2014; Tashiro, Matsubara, and Uehara, 2017). A CVAEs is a generative probabilistic model based on multilayer neural networks. Given an input data *x* and a condition *y*, a CVAE builds a model of the conditional probability log *p*(*x|y*). We used a CVAE as a classifier based on log-likelihood. We emphasize that the CVAE is not based on the 140 × 140 Pearson’s full correlation matrices but is based on 140-dimensional vectors, each corresponding to the intra-regional average signal intensities of the 140 ROIs at one time point. Let 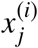 denote the signal intensity of the i-th ROI obtained from a subject at time point *j*. Each sample is a 140-dimensional vector 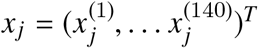 and all the samples obtained from a subject is represented by the set *X* = {*X_j_*}. The CVAE consists of two neural networks called encoder and decoder. The encoder accepts a sample *x_j_* and the condition *y* of a subject, and infers a posterior probability of the latent variable *z_j_*, which is considered to correspond to a nuisance component unrelated to the disease such as something comes into the subject’s mind at the time point *j*. The decoder accepts the latent variable *z_j_* and the condition y, and generates an artificial sample 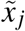. After training the encoder and decoder jointly, the CVAE reconstructs a given signal *x_j_* under the condition *y* accurately, and the reconstruction error indicates (an upper bound of) the negative log-likelihood – log*p*(*x_j_|y*) of the given signal *x_j_* (see the original study (Kingma et al., 2014) for details). Since we considered that each sample *x_j_* was sampled independently from each other, the log-likelihood log *p*(*X|y*) of all the samples *X* of a subject was equal to the summation of the log-likelihood of each sample, i.e., log *p*(*X|y*) = Σ_j_ log *p*(*X_j_|y*). Given the samples *X*, the posterior probability *p*(*y|X*) that the subject belongs to the class *y* is assumed as *p*(*y|X*) = *p*(*X|y*)*p*(*y*)/*p*(*X*) ∝ *p*(*X|y*)*p*(*y*) according to Bayes’ theorem. Therefore, *p*(*X|y* = 1)*p*(*y* = 1) > *p*(*X|y* = 0)*p*(*y* = 0) indicates that the subject is classified into the class *y* = 1. Finally, we assumed that *p*(*y* = 1) = *p*(*y* = 0) = 0.5 for adjusting the imbalance.

We used the encoder and the decoder consisting of 4 layers: The number of units were denoted by *n*_0_, *n*_1_, *n*_2_, and *n*_3_. The hidden layers employed ReLU and layer normalization as their activation functions, and the output layers employed identity function. The condition y was represented by the bias terms of the second hidden layer of the encoder and the first hidden layer of the decoder. The CVAE was trained by Adam optimization algorithm with the parameter *α* = 10^−3^, *β*_1_ = 0.9, and *β*_2_ = 0.999. For each learning iteration, 10 patients and 10 controls were randomly chosen from each site, and 50 samples were randomly chosen from each of the chosen subjects, indicating that a mini-batch comprised 2000 samples. The number of units were searched for within the ranges of *n*_0_ = 140, *n*_1_ ∈ {50,100, 200}, *n*_2_ ∈ {50,100,200}, and *n*_3_ ∈ {5,10} using a Leave-Four-Subjects-Per-Group-Out CV for *validation model 1* and a 10-fold CV for *validation model 2,* where the 10-fold CV was nested within the LTSO CV. We also used *python* 2.7.12 for this analysis.

We can obtain a marginal log-likelihood 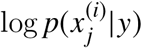 of the *i*-th ROI at the time point *j* using the trained CVAE. A large difference between the marginal log-likelihood 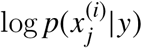 given the different class labels *y* = 0 and *y* = 1 indicates that the *i*-th ROI largely contributes to the classification. Hence, we defined 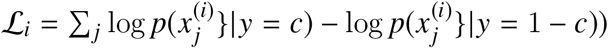 given the correct label *c* as the contribution weight of the *i*-th ROI.

### Characterisation of network changes: Hub disruption

To study the topology of brain networks, we thresholded the Pearson’s full correlation matrices to a produce binary adjacency matrix (consisting of 1’s and 0’s) for each subject. Each of the correlation matrices was thresholded in an adaptive manner to produce an adjacency matrix with a 10 % link density. This value was chosen based on previous studies that have found such a link density to provide optimal discriminative ability (Achard et al., 2012; A. Mansour et al., 2016; Termenon et al., 2016; Itahashi et al., 2014). Using the adjacency matrices, we calculated the Hub Disruption Index (HDI) - a well-recognized method that characterises functional reorganisation in resting-state brain networks in disease (Achard et al., 2012; A. Mansour et al., 2016; Termenon et al., 2016; Itahashi et al., 2014). HDI is calculated based on the difference in a nodal graph-theoretic property of the network, and references the distribution of this metric across all nodes in a single subject, in comparison to the equivalent referential distribution in a cohort of healthy controls. Nodal degree is the most-used index, but any nodal graph measure can be used. Using the Brain Connectivity Toolbox (Rubinov and Sporns, 2010), we examined HDIs for degree, clustering coefficient, betweenness centrality, eigenvector centrality, K-coreness, flow Coefficient, local efficiency, and participation coefficient. This choice is based on the measures that have been applied in previous studies (Achard et al., 2012; Hashmi, Kong, et al., 2014; A. Mansour et al., 2016).

Each of the HDIs, defined as a summary of profile of nodal topological metrics in either a patient compared with the cohort of healthy controls or a control out of the cohort of healthy controls compared with the rest of the cohort of healthy controls (in the same manner as leave-one-out cross-validation), were compared between groups by two-sample two-tailed t-tests and the differences were assigned statistical significance at p values less than 0.05.

### Characterisation of network changes: Modular reorganisation

In order to study the architecture of brain networks in more depth, to identify (and localise, where possible) key differences between patients and control groups, we next probed the network’s modular structure. Our approach was designed to focus on the differences in brain network modularity between patients and control groups by identifying a measure of the consensus modularity pattern across all of the data. We did this by using a new method based on calculation of the multislice modularity (Mucha et al., 2010) - which allows estimation of the basic community or modular architecture across large and complex network data sets. In the *categorical* multislice modularity algorithm (Jeub et al., 2011), the same node is coupled among all subjects of the same group using a coupling parameter ω to create a single symmetric agreement matrix representing each group, see Fig. 1.

**Figure 1:**
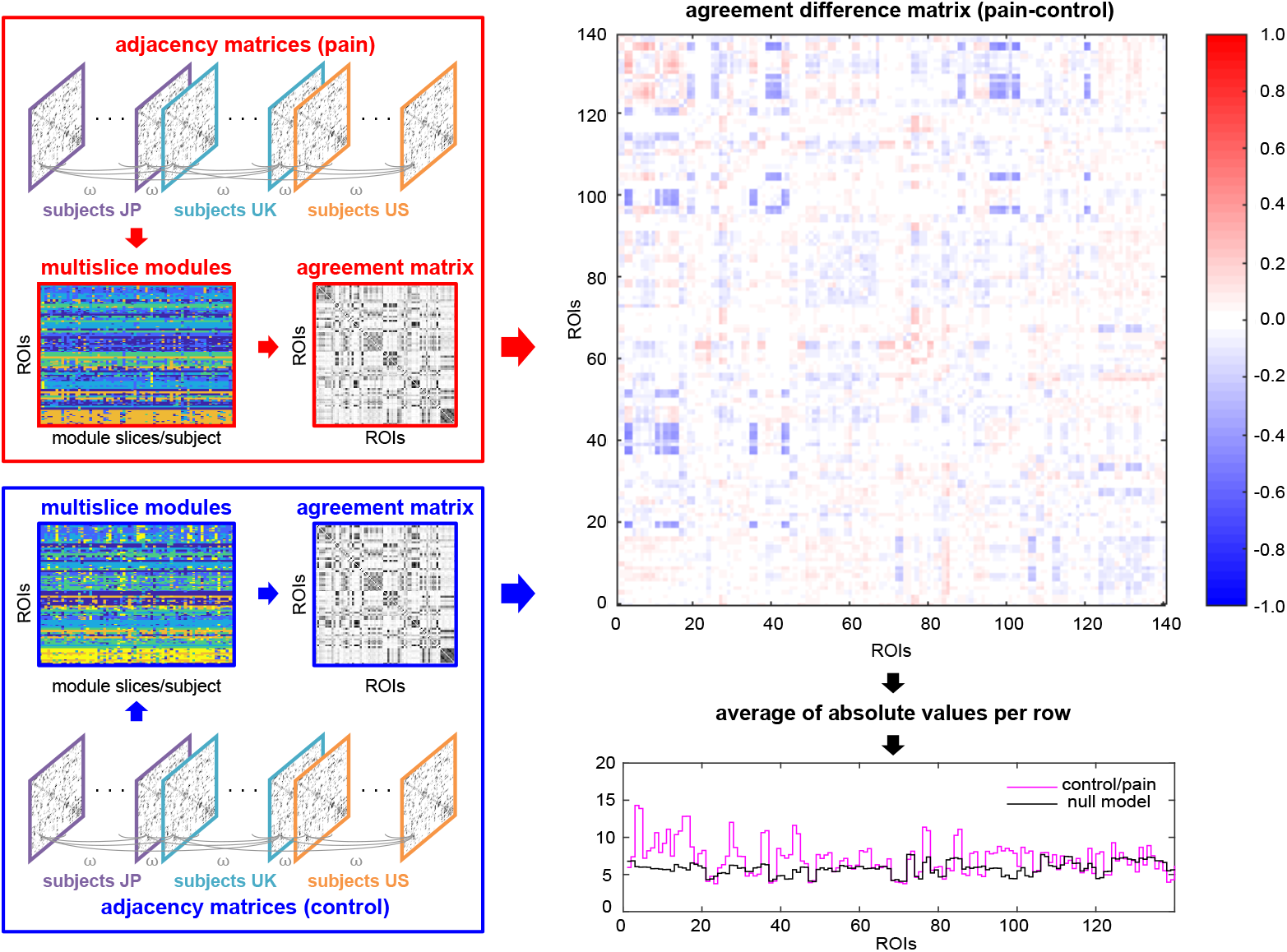
Overview of the computation pipeline for multislice modularity and agreement matrices. First, we calculated the multislice modularity and agreement matrices separately for the pain and control groups, and then calculated their difference. This difference matrix consists of positive values (red) which reflect the likelihood of the two corresponding ROIs (defined by the row and column index) appearing in the same module in the pain group, but not in the control group. Negative values (blue) reflect the opposite - that the two ROIs are less likely to be in the same module in the pain group. Furthermore, values that are near zero (white) reflect pairs of ROIs the do not significantly change or had near-zero agreement in pain and control. The absolute sum of positive and negative values yields an overall metric of modular reorganisation for each ROI (purple plot in lower panel), which can be compared to a chance level calculated from random permutations of the pain and control groups.

This agreement matrix is generated with two free parameters, which we defined *a priori*. First, we chose a modularity resolution of *γ*\* = 1.5, given that this leads to roughly 10-20 modules overall, which is consistent with known architecture of brain networks. Second, we chose a ‘moderate’ coupling strength of *ω*\* = 0.1 based on (Mucha et al., 2010).

The agreement matrix was estimated across all three data sets, separately for the pain patients and the control groups. Since there are slightly more control subjects (*n* = 87) pain patients (*n* = 71), we selected a subset of 70 subjects randomly from each group to match the estimation between each group. Since the modularity estimation is a probabilistic procedure, we repeated this 1000 times, selecting the 70 subjects randomly each time, and computed the average agreement matrix across all repetitions.

We next defined an agreement difference matrix AD as the difference of agreement matrices of pain *minus* that of the control group. Since each agreement matrix has entries within [0,1], large positive entries in AD represent those node pairs that have high agreement in pain, i.e., nodes that are frequently in the same modules for the pain group, but not in the same modules in the control group. Similarly, large negative entries indicate the opposite case, i.e., nodes that frequently join the same modules in the control group, but are not in the same module for the pain group. Nodes with agreement differences near 0 indicate that two nodes are either in the same module for both groups or they are in different modules.

Since the agreement difference matrix has both positive and negative entries in each column, we independently summed the positive-valued and negative-valued elements. This permitted computing a profile of the strongest contributing ROIs in both cases. The sum of the positive and negative contributions provides an overall metric of modular reorganisation for each ROI.

To statistically evaluate the modularity of each ROI, we performed an approximate permutation test, in which we mixed and randomly resampled the pain and control subjects into two groups, and repeated the full analysis. We did this also 1000 times, and calculated the one-sided p-value based on the proportion of times the resampled modularity reorganisation metric exceeded the value based on the correctly specified groups. These results are presented uncorrected for multiple comparisons (across ROIs) below an arbitrary threshold of *p* < 0.01. However we had prior hypotheses related to the 3 sets of regions commonly implicated in chronic pain: sensorimotor cortices, insular-cingulate cortices, and striatal-medial prefrontal cortex.

## Data Availability

### fMRI data

the data from both UK and Japan are available at the ATR Open Access Database https://bicr-resource.atr.jp/decnefpro/ (as raw connectivity matrices).

For the validation fMRI data, we used existing open data from the OpenPain Project (OPP; Principal Investigator: A. Vania Apkarian). OPP funding was provided by the National Institute of Neurological Disorders and Stroke (NINDS) and National Institute of Drug Abuse (NIDA). OPP data are disseminated by the Apkarian Lab, Physiology Department at the Northwestern University, Chicago. Our fMRI data (as dicom files) are in the process of being uploaded here).

### Analysis code

SVM data analysis code for fMRI data (PRoNTO) was co-written by MR is available at http://www.mlnl.cs.ucl.ac.uk/pronto/. The code for modularity reorganisation is available on our github site: https://github.com/leiken26/pain-network. Toolboxes for deep learning analyses are available at https://www.tensorflow.org/about/bib, and we will make openly available user-friendly code for imaging data in due course.

## Results

### Classification using Machine Learning (Support Vector Machine)

Using *validation model 1* (i.e. training on the UK and Japan data and validating on the US data) and correlation as features for classification, the SVM framework correctly predicted 70%, p-value < 10^−3^, of patients (sensitivity) and 56%, p-value < 10^−3^, of controls (specificity), corresponding to a total accuracy of 63%, p-value < 10^−3^. Using *validation model 2* (i.e. training and testing using all available data, UK, Japan and US, with LTSO-CV) and the same features, the SVM framework correctly predicted 68%, p-value < 10^−3^, of patients (sensitivity) and 67%, p-value < 10^−3^ of controls (specificity), corresponding to a total accuracy of 68%, p-value < 10^−3^. The classification results are summarised in Table 2.

**Table 2:**
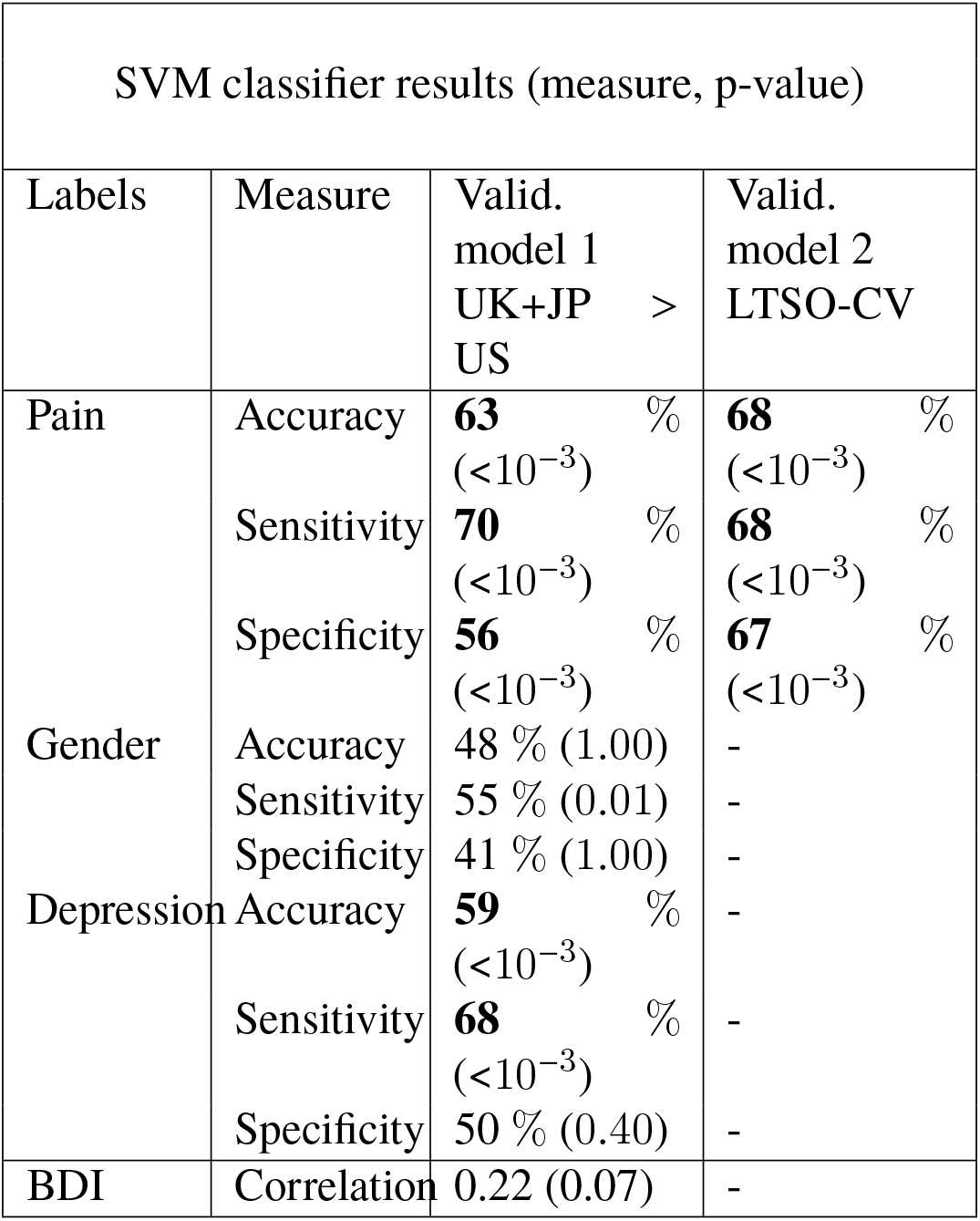
SVM classification results, showing the accuracy, sensitivity and specificity for the two validation models for pain, and also for gender and depression.

To test whether these results were driven by confounds such as gender (the sample was not gender balanced) or depression (many patients also had a depression diagnosis) we tested two new models where instead of the original labels (patient or control) we used ‘male’ and ‘female’ and ‘depressed’ and ‘not-depressed’, respectively. We used exactly the same SVM framework described above and *validation model 1* (trained on UK and Japan data and validated on US data). To obtain the depression-related labels we divided the subjects according to their Beck Depression Inventory (BDI) scores: BDI >= 3 (depressed), BDI < 3 (not-depressed).

The accuracy of the gender model was only 48% (p-value = 1.00) with sensitivity = 55% (p-value = 0.01) and specificity = 41% (p-value = 1.00). The accuracy of the depression model, although statistically significant, was lower than with the pain-related labels: accuracy = 59% (p-value < 10^−3^), sensitivity = 68% (p-value < 10^−3^), specificity = 50% (p-value = 0.40). This result was expected given that the depression labels are highly correlated with the pain labels.

Finally, we also tested if the output of the classifier for each validation sample (i.e. how far is the sample from the decision boundary for both sides) correlated with the BDI score for each individual. The correlation was found to be low (0.22) and not statistically significant (p-value = 0.074). This result together with the two confound models are consistent with the hypothesis that the classifier is related to pain.

### Classification using Deep Neural Networks

Deep learning algorithms represent a novel approach to classification for complex data sets, and have recently been applied to neuroimaging data (J. Kim et al., 2016; Plis et al., 2014; Suk and Shen, 2013). Here, we used a CVAE with the units of *n*_1_ = 100, *n*_2_ = 50, and *n*_3_ = 10, which achieved the best validation accuracy for *validation model 1* (i.e. training on the UK and Japan data and validating on the US data). The CVAE correctly predicted 55% of patients (sensitivity) and 72% of controls (specificity) on average of 100 trials, corresponding to a total accuracy of 63%. Recall that we built a model *p_k_(X|y)* for *k*-th trial. We used the likelihoods Π_*k*_ *p_k_*(*X|y*) of all the 100 models for an ensemble. Then, Π_*k*_*p_k_*(*X|y* = 1)*p*(*y* = 1) > Π_*k*_*p_k_*(*X|y* = 0)*p*(*y* = 0) indicates that the subject is classified into the class *y* = 1. The ensemble achieved a total accuracy of 68%.

Contribution weights varied over trials, and were relatively evenly matched across contributing nodes: Table 5 summarizes the regions which were frequently highly weighted.

**Table 3:** Classification using support vector machine: top 10 positive weights based on validation model 1 (from UK, Japan data, tested on the US data).

| Weight | ROI | ROI Name | MNI Centroid Coords |
| --- | --- | --- | --- |
| 0.0207 | 83 – 128 | R. olfactory sulc. — L. Hippocampus | (12,22,-18) – (-25,-22,-15) |
| 0.0185 | 62 – 72 | R orb. front. sulc. – L. ant. occipito-temporal lat. sulc. | (42,52,0) – (-41,-21,-28) |
| 0.0179 | 42 – 111 | R. cent. sulc. – R. post. inf. temp. sulc. | (42,-17,49) – (54,-60,1) |
| 0.0178 | 51 – 128 | L. ant. inf. frontal sulc. – L. Hippocampus | (-48,39,1) – (-25,-22,-15) |
| 0.0174 | 81 – 138 | R. occipito-polar sulc. – L. cerebellum | (15,-94,-4) – (-25,-61,-35) |
| 0.0164 | 118 – 132 | L. post. branch of sup. temporal sulc. – R. Caud. | (-47,-66,17) – (13,10,10) |
| 0.0163 | 109 – 132 | R. ant. infer. temp. sul. – R. Caudate | (62,-24,-19) – (13,10,10) |
| 0.0162 | 13 – 111 | L. ant. sub-cent. ramus lat. fiss. – R. post. inf. temp. sulc. | (-48,0,7) – (54,-60,1) |
| 0.0155 | 43 – 111 | L. cent. sylvian sulc. – R. post. inf. temp. sulc. | (-60,-2,16) – (54,-60,1) |
| 0.0139 | 61 – 66 | L. orb. front. sulc. – R. sup. frontal sulc. | (-41,50,1) – (26,24,49) |

**Table 4:** Classification using support vector machine: top 10 negative weights based on validation model 1 (from UK, Japan data, tested on the US data).

| Weight | ROI | ROI Name | MNI Centroid Coords |
| --- | --- | --- | --- |
| -0.0192 | 69 – 135 | R. ant. intralingual sulc.. – R. Hippocampus | (15,-60,-4) – (27,-20,-15) |
| -0.0168 | 16 – 62 | R. post. sub-cent. ramus lat. fissure – R. orb. frontal sulc. | (49,-12,16) – (42,52,0) |
| -0.0164 | 8 – 87 | R. asc. ramus of lat. fissure – R. int. parietal sulc. | (50,19,5) – (6,-56,44) |
| -0.0162 | 26 – 61 | R. sup. postcent. intraparietal sulc. – L. orb. frontal sulc. | (48,-28,48) – (-41,50,1) |
| -0.0157 | 106 – 117 | L. rhinal sulc. – R. ant. branch of sup. temporal sulc. | (-26,-7,-36) – (56,-47,27) |
| -0.0154 | 62 – 124 | R. orbital frontal sulc. – L. Thalamus | (42,52,0) – (-10,-19,7) |
| -0.0152 | 1 – 100 | L. ant. lateral fissure — L. sup. precentral sulc. | (-33,13,-21) – (-39,-6,51) |
| -0.014 | 20 – 21 | R. calloso-marginal post. fissure – Left calcarine fissure | (8,-27,45) – (-10,-65,4) |
| -0.0143 | 6 – 62 | R. ant. ramus of lat. fissure – R. orbital frontal sulc. | (45,27,-2) – (42,52,0) |
| -0.0140 | 26 – 42 | R. sup. postcentral intraparietal sulc. – R. central sulc. | (48,-28,48) – (42,-17,49) |

**Table 5:** Classification with deep learning: ROIs which are frequently significant in the CVAE for validation 1

| Frequency | ROI number | ROI name | network | MNI coord. |
| --- | --- | --- | --- | --- |
| 0.116 | 41 | Left central sulcus | sensorimotor | (-41,-20,48) |
| 0.116 | 36 | Right insula | sensorimotor | (42,4,2) |
| 0.116 | 83 | Right olfactory sulcus | default | (12,22,-18) |
| 0.114 | 59 | Left median frontal sulcus | default | (-15,20,58) |
| 0.112 | 44 | Right central sylvian sulcus | sensorimotor | (61,0,17) |
| 0.111 | 23 | Left collateral fissure | default | (-25,-45,-13) |
| 0.111 | 46 | Right subcallosal sulcus | default | (4,-14,25) |
| 0.111 | 40 | Right paracentral lobule central sulcus | cingulo-opercular | (4,-30,55) |
| 0.110 | 33 | Left parieto-occipital fissure | default | (-9,-69,22) |
| 0.110 | 42 | Right central sulcus | sensorimotor | (42,-17,49) |

Using *validation model 2* (i.e. pooling the three datasets together (UK, Japan and US) and using an LTSO CV), the CVAE correctly predicted 56% of patients (sensitivity) and 71% of controls (specificity) on average of 10 trials, corresponding to a total accuracy of 64%. The ensemble of the 10 trials achieved a total accuracy of 68%.

### Characterisation of Network Changes: Hub Disruption

Evidence of reliable network-based classification suggests a possible disturbance of network topology in chronic pain. One way to investigate this further is to apply graph theoretic measures, which allow characteristation of the basic network topology of brain networks (Rubinov and Sporns, 2010). This approach has been widely applied to brain data across a range of psychiatric and neurological conditions (Bullmore and Sporns, 2009; Bressler and Menon, 2010). Of particular relevance is ‘hub disruption’, which refers to a change in the nodal graph topology for any individual metric across the whole brain (Achard et al., 2012). It has previously been shown that brain networks undergo hub disruption for *degree* (the number of connections for each node) in chronic pain, with evidence from both in human chronic back pain patients and rodent pain models (A. Mansour et al., 2016; Termenon et al., 2016). Here, we estimated hub disruption indices across all 3 data sets using a range of nodal graph metrics (see methods). As shown in Fig. 2, we found changes in HDI for clustering coefficient and betweenness centrality consistently across all 3 data sets, and evidence for changes in degree HDI in the US cohort, but not other cohorts.

**Figure 2:**
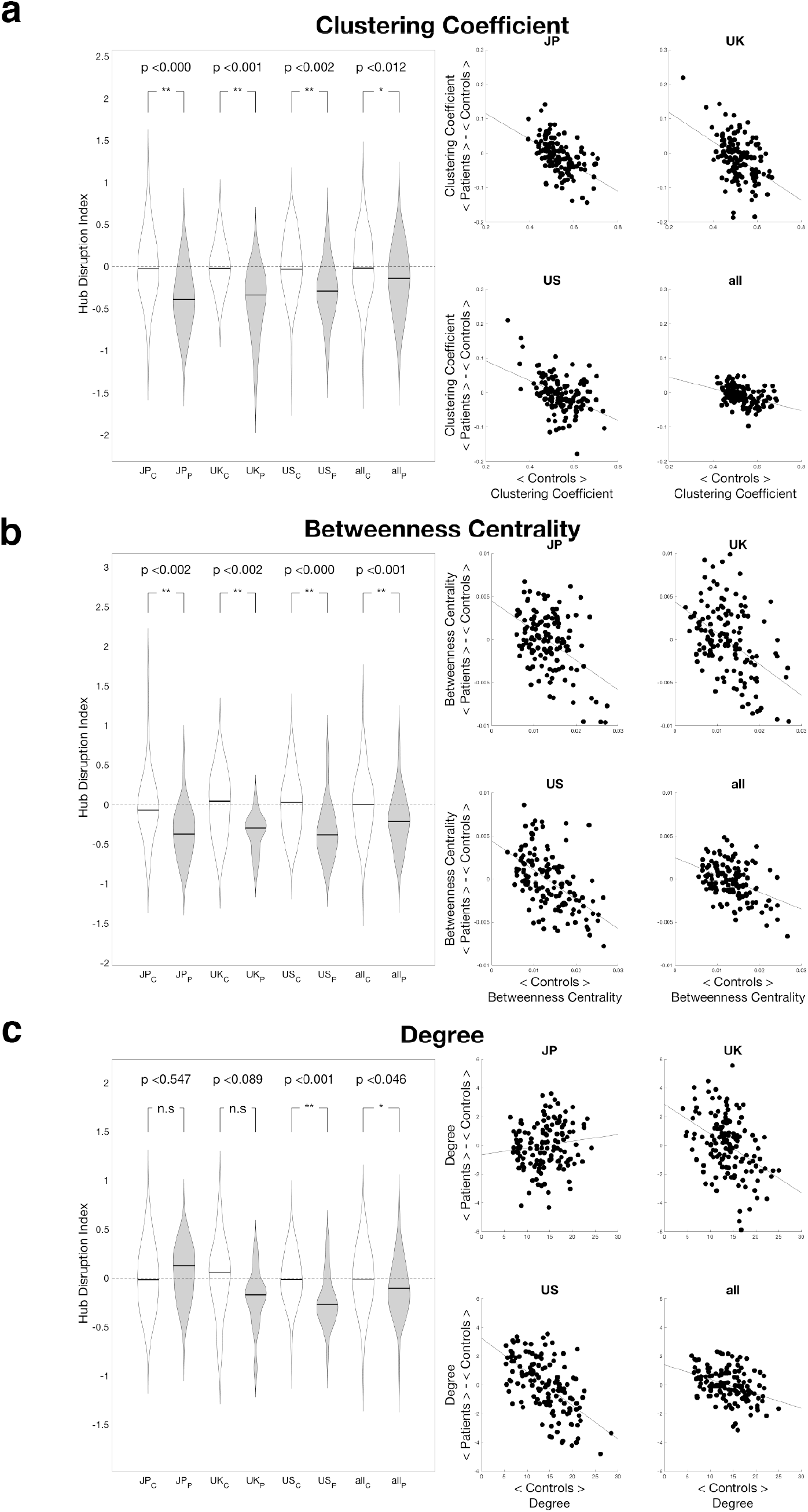
Hub disruption results for a) Clustering coefficient, b) Betweenness centrality, and c) Degree. The figure shows the HDI index individually for each site, and for the entire dataset. For each metric, we show the distribution of subject-wise HDI on the left panels, and the scatter plot of the ROI-specific changes in nodal graph metric on the right panels.

### Characterisation of Network Changes: Modular Reorganisation

The modular structure of the brain - the fact that certain groups of brain regions are especially well-connected with each other, is one of the fundamental properties of brain networks (Meunier, Lambiotte, and Bullmore, 2010; Sporns and Betzel, 2016; Nicolini and Bifone, 2016). Different modules reflect information processing subnetworks that have some degree of independence from each other. The pattern of changes in hub disruption index might suggest a change in the underlying modularity of the network. More specifically, a reduction in the extent to which nodes tend to cluster together, and a reduction in betweenness centrality (the number of shortest paths between nodes that pass through a node), in the absence of other aspects of hub disruption, could relate to a reorganisation of the modular architecture of the network.

To investigate the pattern of modular reorganisation across our chronic pain and control datasets, we developed a method to estimate the common modular architecture across all subjects in each group. Specifically, we computed the multislice modularity within each group, which effectively couples together all subjects within each group into a single large network, and estimates the modular structure of this graph to compute a *consensus* (or ‘*agreement*’) matrix (Lancichinetti and Fortunato, 2012). Then, we computed the difference between the agreement matrix for the chronic pain and control groups, to determine the agreement difference matrix (Fig. 1). This matrix consists of positive (red) and negative (blue) values. The positive values reflect pairs of nodes that are estimated to appear more commonly in the same module in pain patients, and the negative values represent pairs of nodes that are estimated to appear less commonly in pain patients. We defined the overall modular reorganisation for each node as the sum of both positive and negative values for each node (i.e. the sum of each column in the agreement difference matrix). That is, the larger the value, the greater the reorganisation (purple plot in Fig. 1).

To statistically evaluate the amount of reorganisation, we performed a permutation test of reorganisation estimation, to yield one-sided p-values across all ROIs. As illustrated in Table 6 at an uncorrected threshold of p < 0.01, changes were most commonly seen across bilateral sensori-motor regions (Fig 3). We also saw significant changes in right inferolateral prefrontal cortex, and bilateral temporal cortical regions. Looking separately at the positive and negative values related sensorimotor regions, we noted that reorganisation tended to be dominated by negative reorganisation i.e. a reflecting a reduction in the number of modular ‘partners’ of sensorimotor regions in patients.

**Table 6:** Brain ROIs that show modular reorganisation at a cut-off threshold of *p* < 0.01. The table lists the ROI by number (in the BSA-AAL composite atlas), with its corresponding anatomical label, region and MNI coordinates. The overall modularity reorganisation metric AD is listed, included it’s decomposition into positive and negative contributory factors. For anatomy, F=frontal; TP=temporoparietal; FP=frontoparietal; P=parietal

| *AD* (+/-) | p-value | ROI | Anatomical label | Anat. | MNI coord. |
| --- | --- | --- | --- | --- | --- |
| 10.63 (4.23, -6.39) | 0.001 | 8 | R. ascending ramus of the lat. fissure | F | (50,19,5) |
| 9.20 (3.54, -5.66) | 0.001 | 10 | R. diagonal ramus of the lat. fissure | F | (54,17,12) |
| 14.30 (5.29, -9.01) | 0.002 | 3 | L. post. lat. fissure | TP | (-54,-20,10) |
| 12.84 (4.62, -8.22) | 0.002 | 16 | R. post. sub-cent. ramus of the lat. fissure | P | (49,-12,16) |
| 12.02 (5.59, -6.43) | 0.002 | 27 | L. intraparietal sulcus | P | (-30,-66,40) |
| 8.20 (1.52, -6.68) | 0.003 | 98 | L. median precentral sulcus | F | (-20,-15,66) |
| 11.10 (3.67, -7.42) | 0.004 | 11 | L. retrocent. trans. ramus of lat. fissure | P | (-62,-20,23) |
| 12.82 (4.59, -8.24) | 0.004 | 15 | L. post. sub-central ramus of the lat. fissure | TP | (-50,-15,13) |
| 8.50 (1.68, -6.83) | 0.004 | 39 | L. paracentral lobule central sulcus | FP | (-5,-29,57) |
| 11.61 (3.90, -7.71) | 0.004 | 43 | L. central sylvian sulcus | F | (-60,-2,16) |
| 13.88 (5.12, -8.76) | 0.005 | 4 | R. post. lat. fissure | TP | (55,-15,13) |
| 7.67 (1.27, -6.40) | 0.006 | 42 | R. central sulcus | FP | (42,-17,49) |
| 7.84 (1.35, -6.49) | 0.006 | 99 | R. median precentral sulcus | F | (17,-15,68) |
| 8.15 (1.45, -6.70) | 0.006 | 103 | R. sup. postcentral sulcus | P | (26,-39,63) |
| 7.78 (1.34, -6.39) | 0.006 | 120 | L. paracentral sulcus | F | (-6,-16,58) |
| 10.48 (3.20, -7.28) | 0.007 | 44 | R. central sylvian sulcus | FP | (61,0,17) |
| 8.17 (1.46, -6.72) | 0.007 | 96 | L. marginal precentral sulcus | F | (-28,-11,60) |
| 8.52 (1.68, -6.84) | 0.007 | 97 | R. marginal precentral sulcus | F | (27,-8,61) |
| 8.04 (1.43, -6.62) | 0.007 | 121 | R. paracentral sulcus | F | (5,-22,58) |

**Figure 3:**
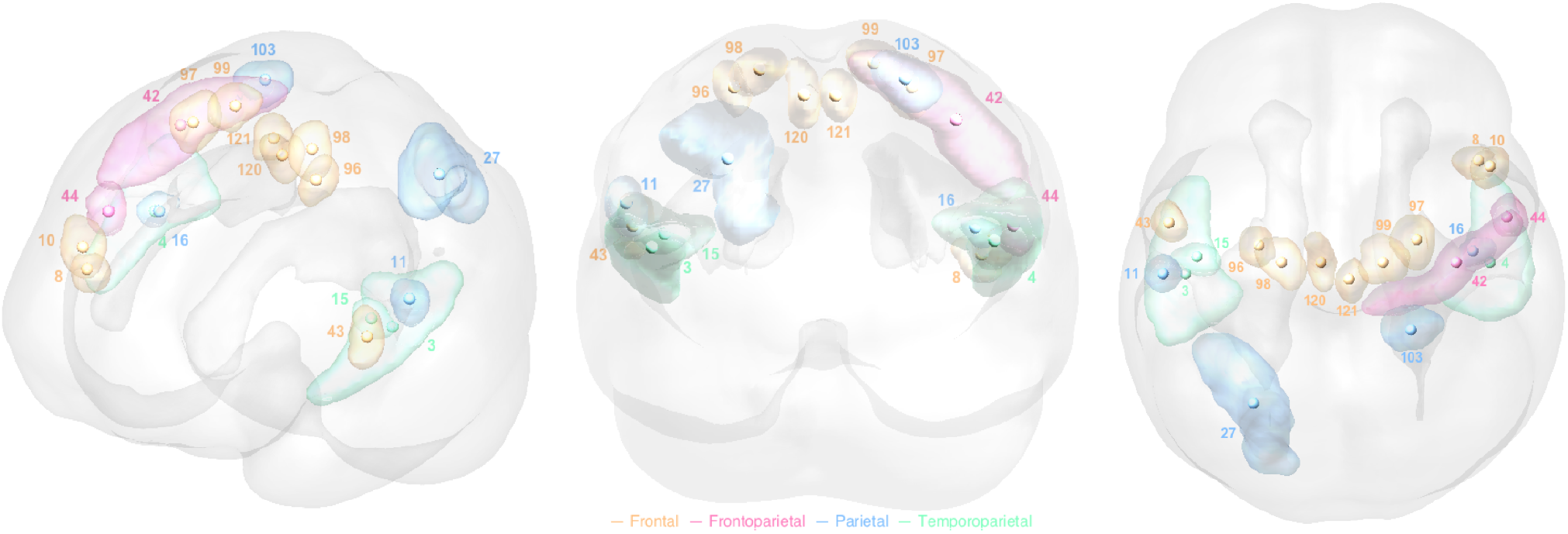
Brain regions showing modular reorganisation. This shows brain ROIs with the best evidence for modular reorganisation in the pain group, compared to the control group, based on the arbitrary theshold of *p* < 0.01, as listed in table 6. The ROIs are colour coded according to their basic anatomical region (cortical lobe).

## Discussion

The results show that there are sufficient brain network changes in chronic back pain to allow reliable classification. Furthermore, the way in which networks are changed follows a characteristic pattern, with global disruption of hub connectivity and modular reorganisation. In particular, we show that bilateral sensorimotor cortical regions undergo the substantial reorganisation, including in regions that also carry predictive weight in classification.

Since chronic pain dominates many aspects of cognition and action, the existence of widespread connectivity changes is not surprising (Baliki, Geha, et al., 2008; Tagliazucchi et al., 2010; Baliki, Baria, and Apkarian, 2011; Kutch et al., 2017; Hemington et al., 2016; Napadow et al., 2010; Hashmi, Baliki, et al., 2013). A challenge, therefore, is to try and identify regions that may have an important or driving role in pain. The approach we take here looks across several methods: connectivity-based machine learning to identify changes important for classification, and a modularity analysis to identify brain regions that show fundamental changes in their functional network identity. Although it is not possible to differentiate causal from consequential connectivity changes, these methods can identify regions that appear to be important in chronic pain at an informational level.

In particular, we present a network modularity analysis approach that aims to identify brain regions that are reorganised in chronic pain. Modular reorganisation is defined on the basis of connections from a particular ROI that appears to join or leave modules with other ROIs - effectively reflecting a change in the ROIs modular identity. This analysis identifies a number of brain regions, but were dominated by sensorimotor cortical regions (i.e. sensory, motor, and premotor cortex). Sensorimotor cortex has been repeatedly implicated in chronic pain (Kutch et al., 2017; Kuner and Flor, 2017; Yanagisawa et al., 2016; Eto et al., 2011; S. K. Kim and Nabekura, 2011; Flor et al., 1997), and consequently is a well-recognised target for intervention. The efficacy of these interventions implies an important role of sensorimotor cortex in chronic pain experience (Yanagisawa et al., 2016; Tsubokawa et al., 1991; Antal et al., 2010; Garcıa-Larrea et al., 1999; Hosomi et al., 2008). Furthermore, voxels in SI have been shown to carry the greatest weight in classifiers trained on BOLD responses to experimental electrical lower back stimulation (Callan et al., 2014), and structural imaging also identify sensory and motor cortices with high classification weights (Ung et al., 2012; Koush et al., 2013), which would be consistent with an important role for multiple subregions of sensorimotor cortex in the chronic back pain state.

Network changes may arise in chronic pain for a variety of distinct reasons, and it is difficult to distinguish these on the basis of a single rsfMRI scan. First, some regions may have a primary causative or risk factor role in the clinical manifestation of chronic pain, and therefore might be expected to be apparent before chronic pain is itself established (e.g. striatal - medial prefrontal cortical regions are candidate regions for this role (Baliki, Petre, et al., 2012). Alternatively, other regions might have no role in the cause or expression of chronic pain, but instead reflect downstream changes, for example perceptual learning of a new sensory environment in which pain is more common (Mancini et al., 2016; Mano, Yoshida, et al., 2017). Such regions might be expected to manifest later, and resolve with successful pain treatment (Rodriguez-Raecke et al., 2009).

Further complications of studying network changes in pain relate to confounding factors such as medication use, secondary effects of pain such as disability, and co-morbid disease such a depression. We also cannot determine the specificity of our results to chronic back pain, as opposed to other chronic pain conditions, and recent evidence suggests that many aspects of network changes may be common (Baliki, A. R. Mansour, et al., 2014; A. Mansour et al., 2016). Hence future network studies can be greatly enhanced by longitudinal data (and pre-morbid data when available), better identifying correlations with pain severity, evaluation of response to drugs, and use of open data sources to provide larger data sets to test generalisation across diagnoses. This should allow identification of components of the network that reflect the cumulative impact of chronic pain, from those reflect a more state-dependent biomarker for ongoing symptomatic experience.

Other caveats that should be noted are that identification of network may depend on the brain atlas used. Higher resolution atlas (i.e. greater number of smaller ROIs) may have a better ability to detection small regions that are important, but greatly increase the numbers of features for classification, which can lead to spurious over-fitting and worse generalisation of the results.

In terms of classification methods, the accuracy of the SVM is comparable with that seen in other disease biomarkers with independent generalisation validation cohorts (Yahata et al., 2016; Takagi et al., 2017) that use machine learning (albeit less than that seen with classifiers for phasic BOLD responses to acute painful stimulation in healthy individuals (Wager et al., 2013)). Here, we also adopted a deep learning approach using deep convolutional neural networks. Deep networks are best known for solving natural image recognition problems with high accuracy, often using very large training data sets. This is typically necessary since they have to extract features from images automatically. With smaller data sets, accuracy is reduced, but performance may still be strong, and this has led to their application to human neuroimaging data. Here, we used a CVAE with a small number of network layers, which suppresses over-fitting in return for a lower classification accuracy than ordinary deep neural networks. An important difference between the connectivity-based decoding and the deep neural network is that the input to the latter is the ROI time-series, not a correlation matrix. In principle, this allows it to use nonlinear and non-pairwise correlations between ROIs implicitly, and hence confers the capacity for much more complex feature extraction. This means that performance may improve when new data becomes available, and help to make deep neural networks a promising method for future classifiers and biomarkers.

In terms of theories of chronic pain, the data here support the general notion of chronic pain as a network disorder, albeit with different aspects of specific regions of the network disturbed in different ways. This approach adds to and complement a substantial body of studies identifying and characterising network changes in chronic pain (Apkarian, Baliki, and Geha, 2009). The limitations of rsfMRI network analysis also also emphasises the importance of understanding the underlying behaviour and computational function of network nodes in chronic pain, and data-driven methods should ideally complement hypothesis-driven task-based studies in clinical groups. Notwithstanding this, a particularly attractive property of the network-theory based approach is their translational applicability to animal models, since topological metrics are relatively independent of brain morphology. In principle, this allows targeted experimental interventions to test whether there is a direct relationship network specific changes and the manifestation of chronic pain.

## Acknowledgements

The authors are grateful to the imaging departments at CiNet (Osaka) and WBIC (Cambridge) for their help with the study.

